# Catalytic bias and redox-driven inactivation of ancestral FeFe hydrogenases from group B2

**DOI:** 10.1101/2023.06.23.541094

**Authors:** Andrea Fasano, Aurore Bailly, Jeremy Wozniak, Vincent Fourmond, Christophe Léger

## Abstract

The biodiversity of hydrogenases, the enzymes that oxidize and produce H_2_, is only just beginning to be explored. Here we use direct electrochemistry to characterize two enzymes from a subgroup of ancestral FeFe hydrogenases, defined by the presence of three adjacent cysteine residues near the active site: the third FeFe hydrogenase from *Clostridium pasteurianum* (CpIII) and the second from *Megasphaera elsdenii* (MeII). To examine the functional role of the unusual TSCCCP motif, which defines the group B2 and is replaced with TSCCP in group A hydrogenases, we also produced a CpIII variant where the supernumerary cysteine is deleted. CpIII and MeII inactivate under oxidative conditions in a manner that is distinct from all other previously characterized hydrogenases from group A. Our results suggest that the supernumerary cysteine allows the previously observed sulfide-independent formation of the Hinact state in these enzymes. We also evidence a second reversible, oxidative inactivation process. Because of their inactivation under oxidative conditions, these enzymes are inefficient H_2_ oxidation catalysts, but their active site itself is not tuned to make them more active in one particular direction.

## Introduction

Hydrogenases, the enzymes that oxidize and produce hydrogen, come in three different flavors -- FeFe-hydrogenases, NiFe-hydrogenases, and Fe-hydrogenases -- named after their metal content. Nitrogenases also produce H_2_ as a byproduct of nitrogen reduction.

The active site “H-cluster” of FeFe-hydrogenases (leftmost in fig 1A) consists of a dinuclear Fe_2_ cluster that is coordinated by 3 CO and 2 CN ligands, a bridging amine (adt, or azadithiolate), and a cysteine thiolate that covalently attaches the dinuclear site to a [4Fe4S] cluster. The Fe ion that is remote from the cubane is referred to as “distal”, or Fe_d_. The diatomic ligands can be detected by FTIR spectroscopy, and their vibrations have been used to identify various states of the H-cluster, including catalytic intermediates^1–5^.

**Figure 1.**
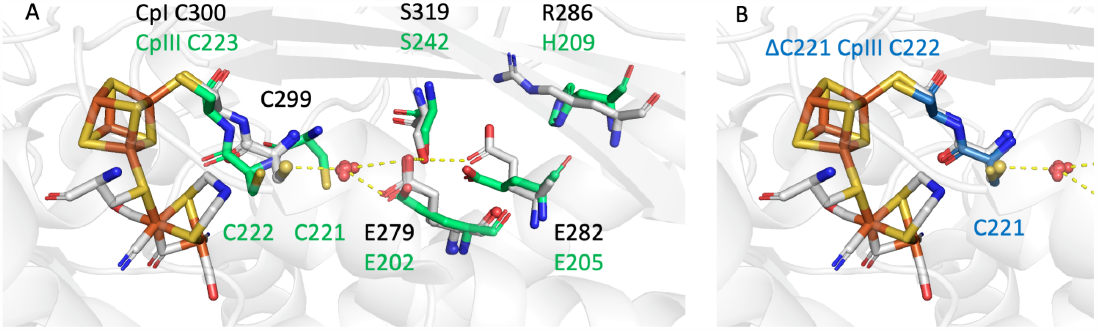
The model structure of CpIII (A) and the ΔC variant (B), aligned on the structure of CpI. Gray sticks indicate the positions of selected CpI residues (light gray, pdb 6N59^5^), the red spheres show the positions of a conserved water molecule in various structures of group A1 FeFe hydrogenases (see the list in Table S1 in ref ^17^). Green sticks show the CCC triad of CpIII and four other residues that are probably involved in proton transfer (2^nd^ rank relaxed AlphaFold model). B: The position of the CC diad in the model structure of the ΔC mutant of CpIII (blue), compared to the position of the CpI residues (gray). SI figure S7 compares the environments of the accessory FeS clusters in CpI and CpIII.

In the catalytic cycle, H_2_ binds to the apical, empty coordination site on Fe_d_ when the active site is in the Hox ‘resting’ state ([4Fe4S]^2+^Fe^II^Fe^I^). H_2_ is split into a hydride and a proton which is transferred to the nitrogen of the amine ligand. The H-cluster is connected to the solvent by a series of residues that mediate long range proton transfer (a conserved cysteine, C299 in figure 1A, is in many hydrogenase the 1^st^ proton acceptor along the chain^3,6^), elusive gaz channels that guide the diffusion of small molecules, and, in some enzymes, accessory FeS clusters that are used to mediate electron transfer.

FeFe-hydrogenases are present in many microorganisms, both prokaryotes and eukaryotes, and are therefore very diverse^7^. The environment of the active site is characterized by three polypeptide motifs (P1, [FILT][ST][SCM]C[CS]P[AGSMIV][FWY], P2 and P3)^8^. Greening et al. have recently defined three phylogenetic groups: A (prototypical and bifurcating), B (ancestral) and C (putative sensory)^9^; Calusisnka *et al* also defined a group D with putative hydrogenases^10^. An earlier, structural classification is based on the number of accessory FeS clusters: zero (M1), two (M2), or four clusters (M3), as exemplified by the three group A1 enzymes from *Chlamydomonas reinhardtii* (Cr), *Desulfovibrio desulfuricans* and *Clostridium pasteurianum* (CpI), respectively. These three standard enzymes have been the focus of most of the biophysical studies of FeFe hydrogenases, but non-standard hydrogenases have recently been characterized and unexpected features have emerged.

For example, the group A “CbA5H” FeFe hydrogenase from *C. beijerinckii* (Cb) is protected from oxygen damage by the binding to the distal Fe ion of the conserved cysteine that is the first proton relay; this binding can occur because non-conserved residues that are remote from the active site make the protein loop that bears the cysteine residue more flexible than in other hydrogenases^4,11,12^. In the group C FeFe hydrogenase from *Thermoanaerobacter mathranii*, protons are transferred along a pathway that is entirely distinct from that of prototypical hydrogenases^13^, and catalysis is ‘irreversible’, that is H_2_ oxidation and production only occur at the price of a large thermodynamic driving force^14^.

Related to the above comment about (ir)reversible catalysis, another catalytic property that has attracted much interest is the ‘catalytic bias’, defined as the ratio of the maximal rates in the two directions of the reaction (H_2_ oxidation and production)^15^. Three homologous FeFe hydrogenases from *C. pasteurianum* illustrate how the protein that embeds the active site can tune the enzyme’s catalytic bias by orders of magnitude: the enzyme CpI is equally efficient in the oxidative and reductive directions, whereas the homologous enzymes CpII (group A) and CpIII (group B) are strongly biased for oxidative and reductive catalysis, respectively^5^.

In 2019, the fact that CpIII is nearly inactive for H_2_ oxidation in solution assays was related by Peters and coworkers to the ease with which the active site is oxidized above its normal resting state Hox. The overoxidized ‘Hox+1’ state is reminiscent of the “Hinact” state observed in other FeFe-hydrogenases, but it is observed in CpIII under less oxidizing conditions. These unique properties were tentatively assigned to the more hydrophobic environment of the H-cluster in CpIII, compared to CpI and CpII, which would destabilize the Hox state and thus favor proton reduction^5^.

At that time, the implication of the observation of the Hinact state in CpIII was not discussed. Only shorty after did it become clear that this state, which is also observed when exogenous sulfide binds to the distal Fe in standard FeFe hydrogenases^16,17^, reveals the binding to the distal Fe of the intrinsic sulfide of a cysteine residue, as occurs in the enzyme from *C. beijerinckii*^*11*^. The hypothesis that this also occurs in CpIII would be particularly appealing because the environment of its H cluster is cysteine rich: the P1 motif, most comonly xTSCCPxx, is present as ITSCCCPMW. This peculiar TSCCCP motif defines the FeFe hydrogenase subgroups ‘M2a’ and ‘B2’ according to Meyer^18^ and Calusinska et al.^10^, respectively. Its functional significance is discussed in this paper, where we examine the catalytic properties of two M2a/B2 enzymes: CpIII (WP_003447632.1) and *Megasphaera elsdenii* II (“MeII”, WP_169013299.1, distinct from the enzyme studied in refs ^19,20^), and a CpIII site directed mutant where the supernumerary cysteine is deleted.

## Results

### Structural considerations

We examined 274 group B sequences listed on the hydrogenase database website^21^ as of may 2023^22^ (we removed WP_025640716.1, which lacks the P1 motif, and included the CpIII and MeII sequences). In this group, the xTSCCPxx motif is the most common version of P1 (134 occurrences), followed by TSCCCPxx (75 occurrences). The latter is specific to group B: none of the 955 hydrogenases sequences classified as group A or C listed on the hydrogenase database included a CCC triad.

In all sequences in group B, the first run of four cysteines, which is Cx2Cx2Cx3C in prototypical hydrogenases, is highly variable (blue dots in SI fig S6), with a consequence on the coordination of the accessory [4Fe4S] clusters that is distal from the active site (SI figure S7), as mentioned before^10,18^.

Figure 1A shows the alphafold^23^ model structure of the WT CpIII, aligned on the structure of CpI (pdb 6N59^5^).The CpI residues are shown in gray. A cluster of red balls indicate the positions of a water molecule that is seen in many structures of group A1 FeFe hydrogenases (see the list in Table S1 in ref ^17^). Proton transfer from the active site to the solvent involves C299, this water molecule (HOH 612 in pdb 3C8Y, absent in pdb 6N59), E279, S319, E282 and R286^3,6^ (black dots in SI fig S6). According to a ClustalO alignment^24^ of 204 group A1 sequences, it is very conserved: the first glutamate is substituted in only 3 sequences, the second glutamate substituted in 4, the serine substituted in only one sequence, and the arginine is substituted (most often with lysine) in only 20 sequences. This series of residues is more variable in group B: the first glutamate is present in all but one group B sequences (WP_013255832.1) out of 274, but the second glutamate is replaced with D or H in about 40 group B sequences (none of which includes the CCC triad) ; the serine is absent from about 30 other group B sequences, and the arginine is rarely present. This shows that there are many small variations on the typical proton transfer pathway in hydrogenases from group B.

The positions of the CpIII residues C223 (equivalent to CpI C300, which coordinates the cubane of the H-cluster), C222, C221, E202, S242, E205 and H209 are shown in green in Figure 1A. The proton transfer pathway is the same in CpI and CpIII, except that a histidine in CpIII replaces R286, and the sulfur of the supernumerary cysteine (C221) is very close to the position of the conserved water molecule. This suggests that C221 mediates proton transfer from C222 to E202 in CpIII. The Alphafold model of MeII (not shown) is similar to that of CpIII, but the putative proton transfer pathway is less well defined ; it would involve C183, C182, E163, E166 but the serine position is occupied by P203 according to the sequence alignment, and the distal arginine by N170 (black dots in SI fig S6).

Figure 1B shows the Alphafold model of the structure of the ΔC221 CpIII mutant where one of the three adjacent cysteines is removed from the sequence: the positions of the two cysteines of the TSCCP motif (blue) match those of CpI (gray), suggesting that the environment of the active site in the ΔC mutant is very similar to that of standard hydrogenases.

### Electrochemistry

Figure 2A compares the voltammetric signatures of Cr, CpIII and MeII FeFe hydrogenases adsorbed onto a rotating pyrolytic graphite edge electrode in a solution saturated with H_2_. In these experiments, electron transfer between the electrode and the enzyme is direct, the electrode potential is swept up and down at a certain scan rate and the activity is measured as a current, positive for H_2_ oxidation, negative for H_2_ production. The electrode is rotated to prevent H_2_ depletion near the electrode surface.

**Figure 2.**
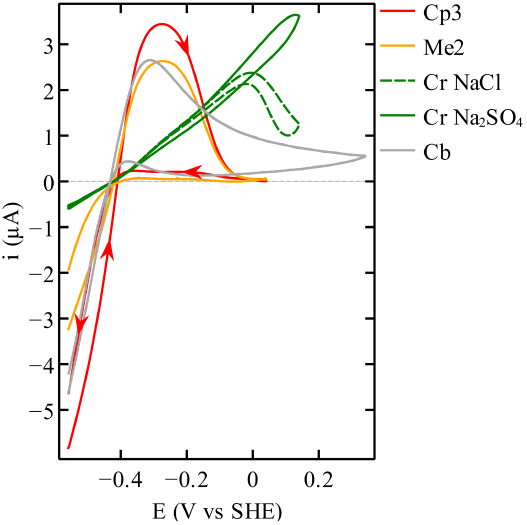
Cyclic voltammetry of CpIII (red), MeII (orange), Cb (gray) and Cr (green), all at pH 7 under one atm. of H_2_. The signals shown using a solid line were recorded without chloride, dashed line in the presence of 0.1M chloride. The Cb voltammogram was scaled down three-fold to ease the comparison. The Cr voltammogram with NaCl was scaled up 1.3 times to compensate for film-loss and allow a proper comparison with the voltammogram with Na_2_SO_4_. Arrows show the directions of the sweeps. A capacitive current recorded in an experiment without enzyme was subtracted from each voltammogram. Conditions: 30°C (except Cb, 5°C); 20 mV/s; 3000 rpm.

The CpIII signal in Fig 2A (red) is consistent with that published in ref ^5^, identical to that of MeII (orange), and very distinct from that of Cr (green) and other standard hydrogenases^25^. The CpIII and MeII enzymes completely inactivate on the scan towards high potentials, above E = -100 mV, but this inactivation is reversible: on the scan towards low potential, a large fraction of the activity is recovered under very reducing conditions (below -400mV): compare the approx. 90% decrease from forward to backward scan at -0.2 V vs the 25% decrease at -0.5 V. SI fig S1 shows the subsequent scan, which demonstrates that the H_2_-oxidation activity is also recovered by the excursion to low potentials. The slight loss of current when the low potential limit of the CV is reached can be due to incomplete reactivation on this particular time scale or film loss.

The CpIII, MeII and Cr CVs shown as plain lines were recorded with the enzyme in a chloride-free buffer at 30°C. Addition of chloride has little effect on the CpIII and MeII CVs (SI fig S2). The reversible oxidative inactivation occurs at a lower potential and is distinct from that of Cr and other group A FeFe hydrogenases : the latter is strictly dependent on the presence of chloride (or bromide) in the buffer^25^, as shown by the dashed line in fig 2. The shape of the CpIII and MeII CVs is reminiscent of that of Cb (gray line in figure 2), although the mechanism appears to be distinct hereafter.

Figures 3A and B show a series of CpIII voltammograms recorded at 5°C (A) and 30°C (B) with the same film and increasing values of the high potential limit (where the sweep is reversed), from red to blue. The enzyme was reactivated by a reductive poise at -509 mV for two to three minutes between each CV.

**Figure 3.**
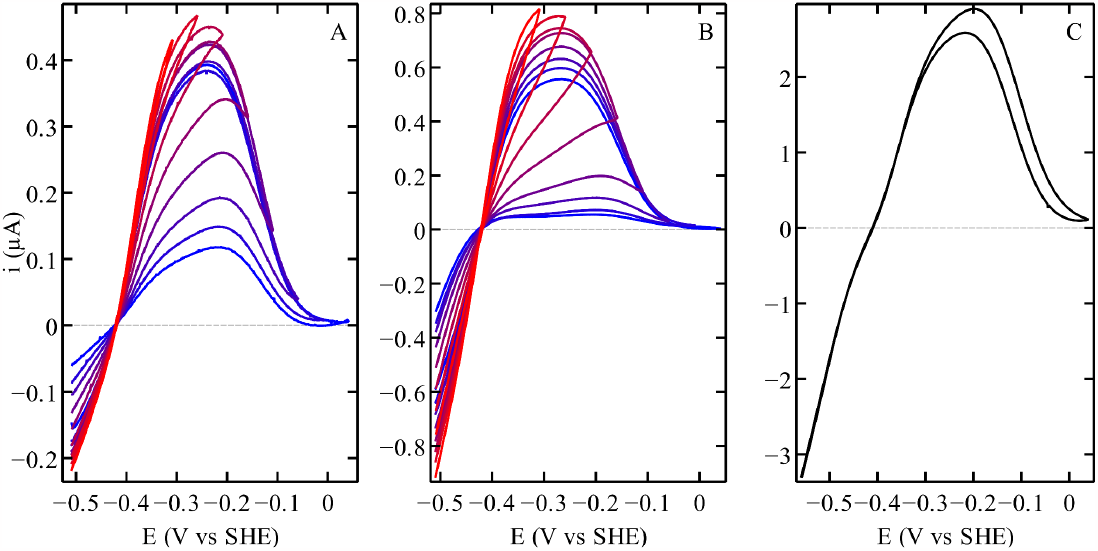
Cyclic voltamograms of WT CpIII recorded by changing the upper potential limits (red to blue from the lower to the greater vertex potential) at 5°C (panel A) and 30°C (panel B). Cyclic voltammogram of the CpIII mutant, missing the supernumerary cysteine of the TSCCCP motif, at 30°C (panel C). The capacitive current recorded in experiments carried out without enzyme was subtracted from all the voltammograms. Other conditions: pH 7; 20 mV/s; 3000 rpm.

The CVs at 5°C in panel A clearly show that the reactivation consists of two different processes. One occurs around -150 mV and results in a sigmoidal increase in current at about the same potential as the inactivation seen on the upward sweep. But this reactivation is not complete: full activity is only recovered at much lower potential, during the reductive poise. This shows that two distinct inactive species, distinguished by their reactivation kinetics, are produced under oxidizing conditions. In the following we call I_1_ and I_2_ the species that reactivate at high and low potential, respectively.

The sigmoidal variations of activity seen around -150 mV, and the observation that the potentials where I_1_ is formed (on the sweep upward) and reactivates (on the sweep backward) are very similar, suggest that I_1_ is in equilibrium with the active form of the enzyme on the time scale of the voltammetry. That this (in)activation is still clearly observed in voltammograms recorded at a much faster scan rate (up to 0.5 V/s in SI fig S5) confirms that it results from a fast reaction. That full reactivation of the enzyme at low potential required a reductive poise (figures 3A and B) shows that the reactivation of the other inactive state, I_2_, is much slower than that of I_1_.

That the sigmoidal reactivation around -150 mV is clearly visible only at 5°C suggests that at high temperature, the formation of I_2_ becomes fast enough that the entire sample is converted into I_2_, a species that reactivates only under very reducing conditions.

The variable high-potential limit experiments in figures 3A and B suggest that I_1_ is not an intermediate between the active form of the enzyme and I_2_. Indeed, if this were to be the case, any increase in the vertex potential (below -150mV) would cause an exponential increase in the rate of formation of I_2_. Instead, the data suggest that the slightly larger amount of I_2_ produced when the vertex potential is increased is the mere consequence of the time spent at high potential being greater. Similarly, it seems possible to rule out the hypothesis that I_2_ is produced from the active form of the enzyme only: in that case, increasing the high potential limit of the sweep would prevent the formation of I_2_, since under these conditions I_1_ would accumulate more. The qualitative inspection of the CVs therefore suggests that I_2_ is produced upon oxidation of either A or I_1_. This hypothesis is supported by the quantitative analysis of the voltammetry below.

We used the following assumptions to fit a model to the CVs in figs 3A and B.

- The active enzyme molecules transform into two fully inactive species, I_1_ and I_2_.

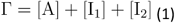

where Γ is the total enzyme surface coverage, and the square brackets indicate surface concentrations (all in units of mol/cm^2^).
- The observed catalytic current *i*(*E,t*) is the product of the steady-state response of the active enzyme times the time-dependent fraction of active enzyme ^26,27^:

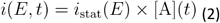
- The steady-state catalytic response of the active enzyme is modeled by the generic “EECr” equation^27,28^.
- I_1_ is in redox equilibrium with the active enzyme.

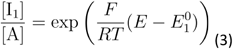

(we expect E°_1_ around -150 mV).
- I_2_ is produced slowly (on the time scale of the voltammetry) either from I_1_ (hypothesis 1), from A (hypothesis 2), or indistinctly from either A or I_1_ (hypothesis 3). If we note *k*_i_ the pseudo 1st-order rate constant of production of I_2_ (inactivation) and *k*_a_ the rate of disappearance of I_2_ (activation), the hypotheses translate into:

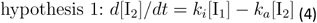

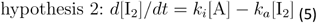

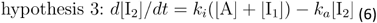
- The rate constants *k*_i_ and *k*_a_ depend on potential in a sigmoidal manner:

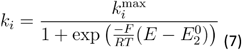

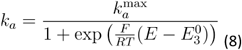

Note that k_i_ tends to k_i_^max^ at high potential, and k_a_ tends to k_a_^max^ at low potential. This (in)activation kinetics is expected if the formation of I_2_ follows an “ECE” mechanism (using the electrochemical terminology), with an oxidation (e.g. the formation of Hox, the most oxidized catalytic intermediate) that precedes the inactivation, a chemical transformation, and a final oxidation that locks the inactive state. But we make no a priori assumption on the values of E^0^_2_ and E^0^_3_.

We used the QSoas software^29^ to fit the model to two successive scans to accurately determine the kinetics of slow reactivation at low potential. Figure 4 shows the best fit of the 3^rd^ model to three selected voltammograms of WT CpIII at pH 7, 30°C (the model was fitted to all the CVs shown in fig 3B globally, we show only 3 CVs in figure 4 for clarity). The other two hypotheses gave less satisfactory modeling (SI figures S3 and S4), as expected from the above qualitative discussion of figure 3. That the 3^rd^ hypothesis gives the best fit of the voltammetry implies that the two inactivation processes are independent of one another. It may be, for example, that one corresponds to a transformation of the active site, and the other results from a remote process.

**Figure 4.**
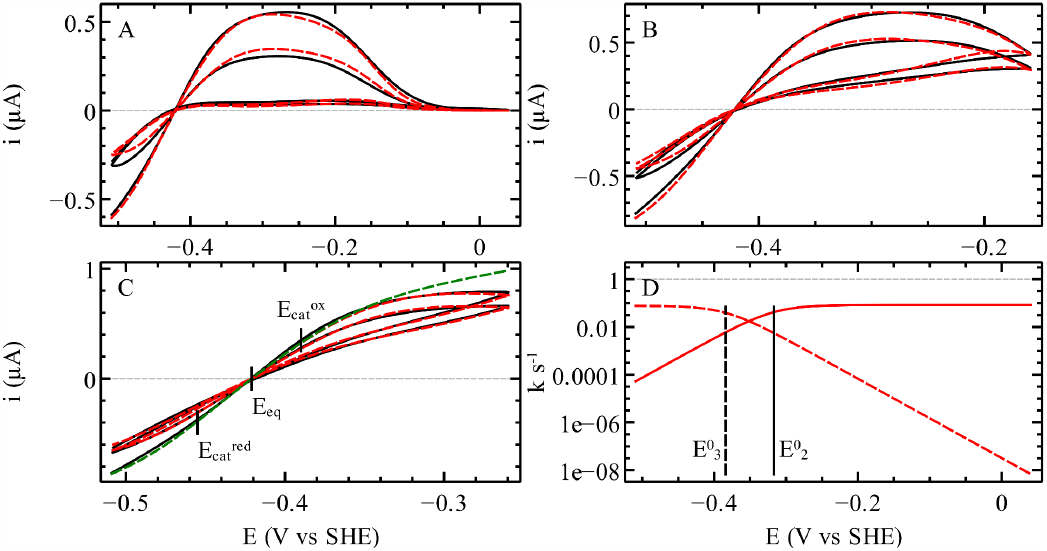
fit of the 3^rd^ model to three selected voltammograms of WT CpIII among those of figure 3B. Panels A, B and C show the experimental data in black and the fits of the 3^rd^ model in dash red lines. Panel D shows the potential dependence of *k*_i_ (solid line) and *k*_a_ (dash line) obtained from the global fit^29^. The green dashed line in panel C shows the steady state catalytic response simulated from the parameters of the best fit, by forcing the inactivation rate constants to be zero. Vertical lines in panel C indicate the two catalytic potentials E_cat_ ^ox^ and E_cat_ ^red^ and the equilibrium potential E_eq_. Solid and dashed vertical lines in panel D note the values of E^0^_2_ and E^0^_3_, respectively.

Panel D shows the dependence of *k*_i_ and *k*_a_ on *E* that is deduced from the global fitting procedure. We obtained k_i_^max^=0.08 s^-1^, k_a_^max^=0.08 s^-1^, E^0^_1_=-139mV, E^0^_2_=-317mV, and E^0^_3_=-384mV for CpIII at 30°C, pH 7. We estimated that the errors on E^0^_2_ and E^0^_3_ are around ± 50 mV. The value of E^0^_1_ defines the equilibrium between the active enzyme and I_1_ at high potential. The value of E^0^_2_ is that of the redox step that precedes the oxidative formation of I_1_; here it is close to the potential of Hred/Hox measured in IR titrations of standard hydrogenases (this parameter has not been measured for CpIII), and the value of E^0^_3_ is close to the value of the reduction potential of the Hox/Hinact transition in CpIII (−380mV at pH 8 in ref ^5^), suggesting that the I_2_ inactive species is Hinact.

In an effort to identify the functional significance of the supernumerary cysteine in the TSCCCP motif, we produced the ΔC221 CpIII variant, by deleting one of the three adjacent cysteine residues. The deletion does not impair activity in either direction, as shown by the CV of the ΔC221 variant in figure 3C. However, the shape of the CV is very different from that of WT CpIII and MeII. In particular, the inactivation seems to involve only the “I_1_” species (which is produced and reactivated at high potential). Indeed, the reactivation at high potential is more pronounced than with the WT and complete. A small hysteresis is observed at high potential; it may also be present in the WT voltammetry but hidden by the second inactivation process (the formation of I_2_), which is not detected in the variant.

In addition to the (in)activation kinetics, the outcome of the fitting procedure is the steady-state response of the fully active enzyme (green in figure 4C) ^27,28^. The latter is defined by two catalytic potentials, *E*_cat_^ox^ and *E*_cat_^red^. The difference between (E_cat_^ox^ + E_cat_^red^)/2 and *E*_eq_, the H^+^/H_2_ Nernst potential, gives the catalytic bias of the enzyme, defined as the ratio of the oxidative and reductive limiting currents, which would be observed in the absence of any inactivation process.

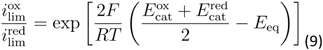

Table 1 shows the values of the catalytic bias of CpIII and Cr, based on the analysis of the voltammograms shown in figures 2A and 4. We conclude that the intrinsic catalytic bias is about the same in all cases, with the enzymes being equally active in both directions of the reaction.

**Table 1.** The catalytic potentials (in mV vs SHE) and catalytic bias under one atm. of H_2_ of Cr and CpIII at 30°C, pH 7, 1 bar H_2_, *E*_eq_ = -419 ± 2 mV.

| | $E_{cat}^{ox}$ | $E_{cat}^{red}$ | $(E_{cat}^{ox} + E_{cat}^{red})/2$ | $i_{lim}^{ox}/i_{lim}^{red}$ | figure |
| --- | --- | --- | --- | --- | --- |
| CpIII | -390 | -455 | -423 | 0.85 | 4 |
| Cr | -362 | -448 | -405 | 2.5 | 2A |

## Discussion

There has been much interest recently in the reaction of the H-cluster of standard^16,17,30^ and non-standard hydrogenases^11,31^ from group A (Cr and Cb, respectively) with extrinsic or intrinsic sulfide. The formation of the sulfide-bound oxidized inactive state is revealed by a particular FTIR signature called H_inact_ and reversed upon reduction. CpIII is another example of hydrogenase that can be observed in the Hinact state^5^, whereas it belongs to the phylogenetically distinct group B. The relations between the formation of Hinact in CpIII, inactivation and protection from O_2_ have not been discussed. The structural determinants of this reaction in CpIII were also unknown, but we have been intrigued by the cysteine-rich TSCCCP motif present in CpIII and other group B FeFe hydrogenases, such as MeII. This motif defines the B2 subgroup.

In the absence of a crystallographic structure of any FeFe hydrogenase from group B, we examined an AlphaFold prediction of the structure of CpIII hydrogenase (Figure 1). The overall fold was already predicted to be very similar to that of standard hydrogenases (SI fig S6 in ref ^5^). The AlphaFold model shown here in figure 1A suggests that the sulfur of the supernumerary cysteine (C222) is involved in proton transfer.

Redox-driven inactivation is a usual feature of FeFe hydrogenases, but significant variations have recently been observed. Oxidative, reversible inactivation is observed with CpI and other standard FeFe hydrogenases from group A, but it occurs only under very oxidizing conditions and only in the presence of chloride, bromide^25^ or sulfide^17^. All experiments herein were performed in a chloride- and sulfide-free buffer, which implies that the mechanism of inactivation is distinct.

Reversible oxidative inactivation occurs with *Clostridium beijerinckii* FeFe hydrogenase (Cb, also from group A, gray in fig 2) in the absence of chloride or exogenous sulfide, and it results from the formation of the Hinact state upon binding to the distal Fe of the H-cluster of the sulfur of the cysteine that is equivalent to CpI C299^11,31,32^ (see figure 1A). The observation that CpIII is also oxidized to the Hinact state in the absence of sulfide (Table S3 in ref ^5^) strongly suggests that the binding of the cysteine to Fe_d_ occurs in CpIII, as it does in Cb. This is probably one of the two reasons CpIII inactivates at high potential. The slow reduction of Hinact may be the cause of the lag observed in solution assays of CpIII (SI section 1.2 in ref ^5^). The voltammetry of CpIII and MeII being very similar, it is tempting to suggest that this bond formation also occurs in MeII and other hydrogenases from group B. The oxidative formation of a vicinal disulfide bridge^33^ between C222 and C221 may also result in enzyme inactivation, but this alternative hypothesis does not explain the observation of the Hinact state, so we considered it unlikely.

The voltammetric signatures of CpIII and MeII in figure 2 may look similar to that of Cb at first (figure 2). However we observed that the (in)activation kinetics of CpIII and MeII is different from that of Cb. In particular two distinct inactive species, referred to here as I_1_ and I_2_, are produced independently.

We consider most likely that the 2^nd^ inactive species (I_2_) is the Hinact state. Indeed, the reduction of Hinact occurs at low potential (−400mV at pH 8 according to the results in ref ^5^), which matches the reactivation kinetics of I_2_ (E^0^_3_=-384mV at pH 7, according to the above described modeling of the voltammetry).

The observation that the binding of the cysteine and the formation of Hinact occur in these group B enzymes, which are phylogenetically distinct from Cb, is intriguing. The three residues that have been identified as allowing the formation of Hinact in Cb by making the TSCCP loop flexible (L364, P386 and A561)^34^ are not conserved in CpIII and MeII (see the orange frames in SI fig S6), but we observed that in CpIII, this inactivation disappears when the supernumerary cysteine is deleted. This suggests that in WT CpIII, C221 pushes the proximal cysteine (C222) closer to Fed and hence favors the formation of the Fed-Scys bond. It therefore appears that different peculiar structural features in Cb on one side, and group B2 hydrogenases on the other, give the protein the ability to allow the proximal cysteine to bind Fe_d_.

Binding of a nearby residue to Fe_d_ should protect the H-cluster from O_2_. This is so in Cb, but the same protection mechanism operates in a certain type of NiFe hydrogenases, where the binding to a metal ion of the active site of a nearby aspartate protects the enzyme from oxygen attack^35,36^. However, we have found no evidence that CpIII hydrogenase is resistant to O_2_. This may be because the formation of Hinact is too slow (0.08s^-1^ at 30°C in CpIII, compared to 1s^-1^ at 5°C in Cb ^11^), so that the disruption of the active site by O_2_ occurs faster than the enzyme can protect itself by forming the H_inact_ state. The relation between fast formation of Hinact and greater resistance was demonstrated in a series of Cb site directed mutants ^11,12^.

The other inactive species, **I**_**1**_, is produced and reduced more quickly and at much higher potential (−150mV) than I_2_, irrespective of the presence of the third cysteine in the P1 motif (that is, in WT CpIII and in the ΔC variant, fig 3C). Since the formation of I_2_ occurs independently from the production of I_1_ at the active site (hypothesis 3 in the modeling of the voltammetry), this inactivation, which is not affected by the change in coordination of Fe_d_, must be the consequence of a remote transformation; we speculate that this is the overoxidation of an accessory cluster, maybe the distal cluster whose coordination pattern is very peculiar in group B hydrogenases (SI figure S7). This hypothesis will have to be tested by mutagenesis.

To discuss the catalytic bias and the shape of the voltammograms, one needs to acknowledge that the catalytic response of the enzyme is modulated by the inactivation processes: the observed voltammogram is actually the product of the steady-state response of the active enzyme (an intrinsic property of the active site) times the time-dependent fraction of enzyme that is active (considering the two redox-dependent, reversible inactivation processes, eq 2). The quantitative modeling of the voltammetry (figure 4) allows us to untangle the two contributions, to obtain the steady-state response (green in figure 4C) and the kinetics of inactivation. The shape of the former tells whether the enzyme is an equally good catalyst in the two directions of the reaction, or if it is biased in one particular direction. This is quantified by the difference between the average catalytic potential and the Nernst potential of the H^+^/H_2_ couple (eq 9). Table 1 shows that CpIII hydrogenase is actually not intrinsically biased in any direction (the ratio of the limiting currents is close to one), just like Cr. This contrasts with the observation that in solution assays^5^, CpIII is much more active in the reductive than in the oxidative direction. The reason for this apparent discrepancy is that in solution assays, the enzyme inactivates under the oxidative conditions that are required to drive H_2_ oxidation, hence the very small oxidative activity. The same is true for Cb, which is very efficient at oxidizing H_2_, but whose strong H_2_-oxidation activity is hidden by the oxidative inactivation, except in some variants where this inactivation is slowed (figure 3d in ref ^11^).

If one is interested in using the enzyme to catalyze H_2_ oxidation, that it inactivates under oxidative condition is clearly an issue that has to be considered. However, if one’s goal is to elucidate how the residues that surround the H-cluster may tune its catalytic properties and the catalytic bias, then any redox driven inactivation must be factored out. Here, we conclude that the properties of the active site of CpIII hydrogenase are not very different from those of group A hydrogenases (e.g. Cb or Cr), in that the enzyme is intrinsically equally active in both directions. That Cb and CpIII hydrogenase inactivate at high potential does not reveal any difference in terms of H-cluster redox and catalytic properties.

## Supporting Information

Molecular biology and biochemistry procedures, additional electrochemical experiments and data analyses, sequence alignments. The SI file is available free of charge at https://pubs.acs.org

## Acknowledgements

This research was funded by the Centre National de la Recherche Scientifique, Aix Marseille Université, Agence Nationale de la Recherche (ANR-21-21-CE50-0041), Région Sud. This work received support from the french government under the France 2030 investment plan, as part of the Initiative d’Excellence d’Aix-Marseille Université – A*MIDEX, AMX-22-RE-AB-097. The authors are very grateful to Frédérique Berger, glass blower at Aix Marseille University and the proteomic facility of the Institut de Microbiologie de la Méditerranée (IMM, CNRS-AMU, FR3479), Marseille Proteomique (MaP), for performing proteomic analyses by mass spectrometry.

