## Supporting information for "Catalytic bias and redox-driven inactivation of ancestral FeFe hydrogenases from group B2"

#### Supplementary information

Andrea Fasano, Aurore Jacq-Bailly, Jeremy Wozniak, Vincent Fourmond, Christophe Léger

Laboratoire de Bioénergétique et Ingénierie des Protéines, CNRS, Aix Marseille Université, Marseille, France.

**Biochemistry** **1**
Gene synthesis 1
Sequences 2
Strains 2
Plasmids 3
Primers used to generate the ΔC221 variant of CpIII 4
Mutagenesis of the C221 deletion 4
Culture 4
Purification 4
**Electrochemistry** **5**
Material and methods 5
Data 5
**Structural considerations** **8**
Sequence alignments 8
Alphafold calculations 10
The coordination of the accessory clusters in CpIII 10
**References** **11**

### Biochemistry

#### Gene synthesis

The protein sequences of the CpIII and MeII hydrogenases characterized in this work were from the NCBI database, accession numbers WP_003447632.1 and WP_036203142.

The DNA sequences (table S1) were optimized for production in E. coli with the softwares Rare Codon Analysis - Online Calculation and Plot Tool - BiologicsCorp and Repeats Finder for DNA/Protein Sequences. The sequences were synthesized by GeneCust and inserted in the MCS1 of pET-Duet1 between NcoI and NotI with a C terminal StrepTag, generating the two plasmids pEt-duet-hydCpIII and pET-Duet-MeII. Each of these plasmids was co-transformed in

E. coli cells BL21(DE3) Δ*iscR* with pACYC-hydGX-EF, expressing the H-cluster maturases HydE, HydG, HydF and HyDX[1], as described for CpIII in ref [2].

### Sequences

Table S1: optimized sequences used in this work for Cp HydIII and Me HydII. The C terminal strap tag is highlighted in red.

Cp HydIII strep tag

ATGAAAAGCGAATATAACGATCTCTTTAAGAGCCTGATTGATGCGTATTATAAAGATGATTTTGATGAAT
TTATTAAAAAAGCACTGAGCGATAGCACCGTGAACAAAGAAGAACTGAGCAACATTATTAGCAGCTTTTG
CGGCGTGGAACTGAAATATACCGATAAAGATACCTATATTAAAGACCTGAAAAACGCGATTAAAAACTAT
AACAGCGATCATAAAATTGTGACCAAAATTCGCGATTGCAGCGTGGATTGCGCGGATGAAAACGGCAAAA
CTAGCTGCCAGAAAAGCTGCCCGTTTGATGCGATTCTGATTGATGAAGCGAACAAAACCAGCTATATTGA
TAAAGACCTTTGCACCGATTGCGGCTTTTGCGTGGAAGGCTGCCCGAACGGCAGCATTCTGGATAAAGTG
GAATTTATTCCGCTGGCGAACCTGCTGAAAGAAAAGCAGCCGGTGATCGCGGCAGTGGCGCCGGCGATTA
CCGGCCAGTTTGGCGATGATGTGACCATTGATCAGCTGCGCACCGCGTTTAAAAAAGTGGGCTTTGCAGA
TATGATTGAAGTGGCGTTTTTTGCGGATATGCTGACCCTGAAAGAAGCGTGCGAATTTAACGCGCATGTG
AAAAGCAAAGATGATCTGATGATTACCAGCTGCTGCTGCCCGATGTGGGTGGGCATGCTGAAACGCGTGT
ATAAAGATATGGTGAAATATGTTAGCCCGAGCGTGAGTCCGATGATTGCGGCGGGCCGCGTGATTAAAAC
CCTGAACAGCAACTGCAAAGTGGTGTTTGTGGGCCCGTGCATTGCGAAAAAAGCGGAAAGCAAAAACAAA
GATATTGAAGGCGATATTGATTTTGTGCTGACCTTTGAAGAAGTGAAAAACATTTTTGAAAGCCTGAATA
TCAACCCGAGCGAACTGCCGGAAGATCCGAGCACCGATTATGCGAGCCGCGAAGGCCGCCTGTATGCGCG
CACCGGCGGCGTGAGCATTAGCGTGAGCGAAGCGGTGGCGAAACTGTTTCCGGAAAAGAAAGACCTGTTT
AAAAGCGTGCAGGCGAACGGCGTGATTGAATGCAAAAAGATTCTGGAAAAAGCGCAGAACGGCGAAGTGG
CGGCGAACTTTATCGAAGGTATGGGTTGTGTGGGCGGCTGCGTGGGCGGCCCGAAAGCGCTGATTCCGAA
AGAAAAAGGCCGCGAAAAAGTGAACGAATTTGCGGAAAACAGCAACGTGAAAATTAGCCTGGAAAGCGAT
CAGATGAAAAAAATTCTGAACATGCTGAACATTACCAGCGCGAAAGATTTTATGGATGAAGAAAAAATTA
AAATTTTTGAACGCGAATTTGCTAGC**TGGTCCCACCCGCAGTTCGAAAAG**TAA

Me HydII strep tag

ATGAAAACCCTGGATGAACTGTACCAGCAGATGCTGAAACGCGCGGCGGAAGGCAAACCGGTGGGCGTGG
ATGGCGATCCGAAACTGATTGATTGCCTGGCGCATCCGGATGATCATCCGCTGATTTGGCGCGTGAAACC
ATGCCTGTGTCCGCCGGATGAACCGCATCATTGCCAGGAAGCGTGCGATTGGGGCGCGATTACCAGCGGC
CCGGATGGCATTGCGATTGATGATAGCCAGTGCGTGGGCTGCCAGGCGTGCGTGGATGCGTGCAAACTGG
ATACCCTGAAAACGCGTAAAGATCTGATTCCGGTGGTGCAAGAACTGCATGATTATGATGGCCCGATTTA
TGCGATGGTGGCGCCGGCGGTGAGCGGCCAGTTTGGCCCGGATATTACCATGGGCAAACTGCGCACCGCG
TTTAAACGCCTGGGCTTTACCGGCATGCTGGAAGTGGCGGTGTTTGCGGATATTCTGACCCTGAAAGAAG
CGCTGGAATTTGATAAAAACATTAACAGCGAAGATCAGTTTCAGCTGACCAGTTGCTGTTGCCCGATGTG
GATTGCCATGATCAAAAAACTGTATCATCAGCTGCTGCCGAACGTGCCGGGCAGCGTGAGCCCGATGATC
GCGGGTGGCCGCACCGTGAAAGCGCTGCATCCGAAAGCGAAAACCGTGTTTATTGGCCCGTGCCTGGCGA
AAAAAGCGGAACGCAAAGAACCGGATCTGGTGGGCGCGGTGGATTATGTGCTGACCTATCAGGAACTGCG
CGATCTGTTTAGCGTGACCAACATTGATCTGGCGAGCCTGCCGGAAGATAACCTGCCGCATGCGAGCGAA
GCGGGCATTAGCTATGCGTATGCGGGCGGCGTGAGTGAAGCAGTGACCGAAACCGTGCATCAGCTGAACC
CGGATCGCCAGGTGGATATTAAAACCCGCAAAGCGGATGGCGTGCCGGCGTGCAAAGCGATGATTAACGA
TATTCTGGCGGGCAACCGCGATGGCAACTTTTTTGAAGGCATGGGCTGCATGGGCGGCTGCGTGGGCGGC
CCGCGCGCGATTATTAAAAAAGAAGAAGGCAAAGCGAACGTGCAGGCGTATGGCAAACAGGCGGCGTATA
AAACCCCGATGGAAAACCCGTATGTGATTGAACTGCTGAAAAAACTGGGCTTTCCGACCGTGGAAGAATT
TCTGGAACGCAGCCATCTGTATGATCGCAAATTTGATGCTAGC**TGGTCCCACCCGCAGTTCGAAAAG**TAA

### Strains

Table S2: the E. coli strains used in this work.

| name | relevant characteristics | Ref | Justification |
|---|---|---|---|

| | | | |
|---|---|---|---|
| *E. coli* DH5α | F - Φ80 *lac ZΔM15* Δ( *lac ZYA*- *arg F)U169* *rec A1* end *A1 hsdR17* (rK–, mK+) *pho A sup E44 thi-1 gyr A96 rel A1 ton A* | New England Biolabs | used for cloning |
| *E.coli* BL21 (DE3) Δ*iscR* | F− ompT hsdSB(rB − mB − ) gal dcm iscR::kan (DE3) | [3] | used for protein production |
| *E.coli* BL21 (DE3) Δ*iscR/*pET-Duet1-*hyd-CpIII/*pACYC-*hydGX-EF* | *E.coli* BL21 (DE3) Δ*iscR* carrying the plasmid pET-Duet1-*hyd_CpIII* and pACYC-*hydGX-EF* | This work | used for CpIII production |
| *E.coli* BL21 (DE3) Δ*iscR/*pET-Duet1-*hyd_CpIII-delC221/*pACYC-*hydGX-EF* | *E.coli* BL21 (DE3) Δ*iscR* carrying the plasmid pET-Duet1-*hyd-CpIII-delC221* and pACYC-*hydGX-EF* | This work | used for CpIII-delC221 production |
| *E. coli* BL21 (DE3) Δ*iscR/*pET-Duet1-*hyd_MeII/*pACYC-*hydGX-EF* | *E.coli* BL21 (DE3) Δ*iscR* carrying the plasmid pET-Duet1-*hyd_MeII* and pACYC-*hydGX-EF* | This work | used for MeII production |

#### Plasmids

Table S3: the plasmids used in this work.

| name | relevant characteristics | Ref | Justification |
|---|---|---|---|
| pET-Duet1-*hyd-CpIII*-streptag | promoter T7, Ampicilin resistance | this work | for CpIII production |
| pET-Duet1-*hyd-CpIII-delC221*-streptag | promoter T7, Ampicilin resistance | this work | for CpIII_delC221 production |
| pET-Duet1-*hyd-MeII*-streptag | promoter T7, Ampicilin resistance | this work | for MeII production |
| pACYC-*hydGX-EF* | promoter T7, resistance to chloramphenicol | [1] | for production maturase of *Shewanella oneidensis* |

#### Primers used to generate the ΔC221 variant of CpIII

Table S4: the primers used in this work.

| | |
|---|---|
| CpIII ΔC221 forward | 5′ ATGATTACCAGCTGCTGCCCGATGTGGGTGGGC 3′ |
| CpIII ΔC221 reverse | 5′GCC CAC CCA CAT CGG GCA GCA GCT GGT AAT CAT 3′ |

#### Mutagenesis of the C221 deletion

The QuikChange II Site-Directed Mutagenesis Kit (Agilent) was used to generate point mutations in the gene of the small subunit of CpIII, *hyd-CpIII*, following the manufacturer's instructions. Oligonucleotides "CpIII ΔC221 forward" and "CpIII ΔC221 reverse" (Table S2) containing the mutation ΔC221, each complementary to opposite strands of the vector pET-Duet1-hyd-CpIII-streptag, were used to amplified this plasmid by PCR. After DpnI treatment, the resulting vector pET-Duet1-*hyd-CpIII-delC221*-streptag was transformed into the *E.coli* BL21 (DE3) Δ*iscR* strain [4] (Table S2).

#### Culture

Culture, growth and purification of CpIII, MeII and CpIII ΔC221 were performed in the same manner. The transformed cells of E.coli BL21 (DE3) ΔiscR were first grown overnight in 10 mL of TB medium supplemented with trace elements, ampicillin (100μM) and kanamycin (50μM) at pH 7, 37 °C, under agitation of 160 rpm. 1 mL of the overnight pre-culture was inoculated in 1 L flasks of TB medium supplemented as before, and let grown aerobically (160 rpm) at 37° C until $DO_{600nm}$ of 0.5. The cultures were then placed in anaerobic condition, at 20°C, and protein expression was induced with 0.5 mM IPTG for 16 hours.

#### Purification

*E.coli* cells were harvested aerobically by centrifugation at 8000g for 20 min at 4° C.
CpIII, MeII and CpIII ΔC221 were purified anaerobically and all the following steps were performed under a nitrogen atmosphere. The pellets were washed with buffer A (Tris-HCl 0.1M pH 8.0, NaCl 300 mM, 2 mM DT).
The pellets were resuspended in 7 mL of buffer per g of cells, using buffer A supplemented with protease inhibitors (SIGMAFAST™ Protease Inhibitor CocKtail Tablet, Sigma-Aldrich) and DNAse I (ROCHE).
The cells were disrupted by 2 passages through an Emusiflex C5 between 10000-15000 psi. We incubated the extract with avidin 2 mg/L for one hour. An ultracentrifugation was performed at 245000 g, at 4°C for 45 min. The soluble fraction was loaded on a resin (2 mL) of StrapTactin-Superflow (IBA). The resin was washed with 20 mL of buffer A and the hydrogenases were eluted by adding 6mL (6 fractions of 1 mL) of Tris-HCl 0.1M pH 8.0, NaCl 300mM, desthiobiotin 2.5 mM and 5% (w/v) glycerol. The concentration of protein was determined with the Bradford assay (Bio-Rad).

### Electrochemistry

#### Material and methods

Cyclic voltammetry experiments were performed in a thermostated electrochemical cell in mixed buffer consisting of MES, CHES, HEPES, TAPS, Na acetate (each 5 mM), and $Na_2SO_4$ (0.1 M). The buffer contained no chloride unless stated otherwise. A saturated calomel was used as reference electrode and a correction factor of +241 mV was applied for conversion to SHE.  All the experiments were run in a glovebox under nitrogen atmosphere. Between every cyclic voltammogram shown in figure 3A and B of the main text, the potential was kept at the value of the lower vertex (-509 mV vs SHE) to allow complete reactivation of the enzyme film before the next experiment. Cyclic voltammetry experiments were recorded with the software GPES and analyzed with the free software QSOAS[5] (www.qsoas.org).

The film of enzymes were performed by drop casting between 0.5 and 1 µL of enzyme solution (between 5 and 10 µM) on the the surface of an home made pyrolytic graphite edge electrode (3 mm diameter) previously polished with alumina powder (Buehler, 1 µm).

#### Data

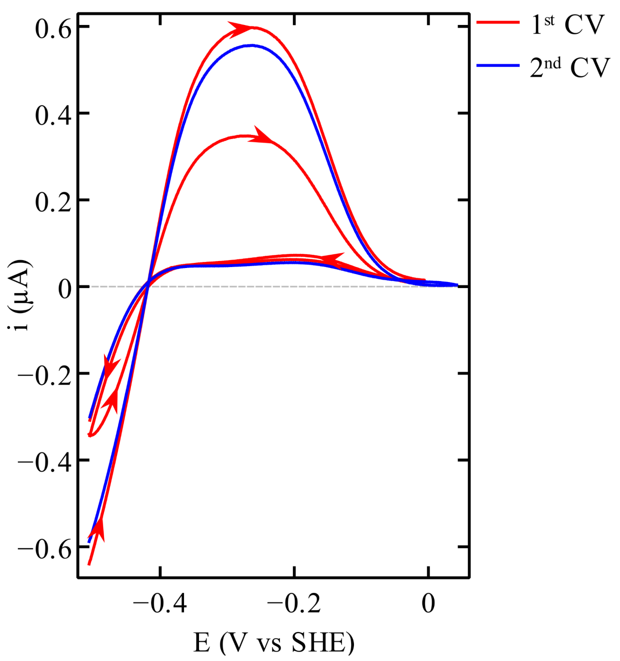


Figure S1: In red two successive scans with CpIII WT. In blue the successive CV performed with the same film of enzyme. Between the two CVs the potential was fixed at -0.509 mV vs SHE, to allow the reactivation. The capacitive current recorded without enzyme was subtracted from all the voltammograms. Conditions: 30°C; pH 7; 20 mV/s; 3000 rpm.

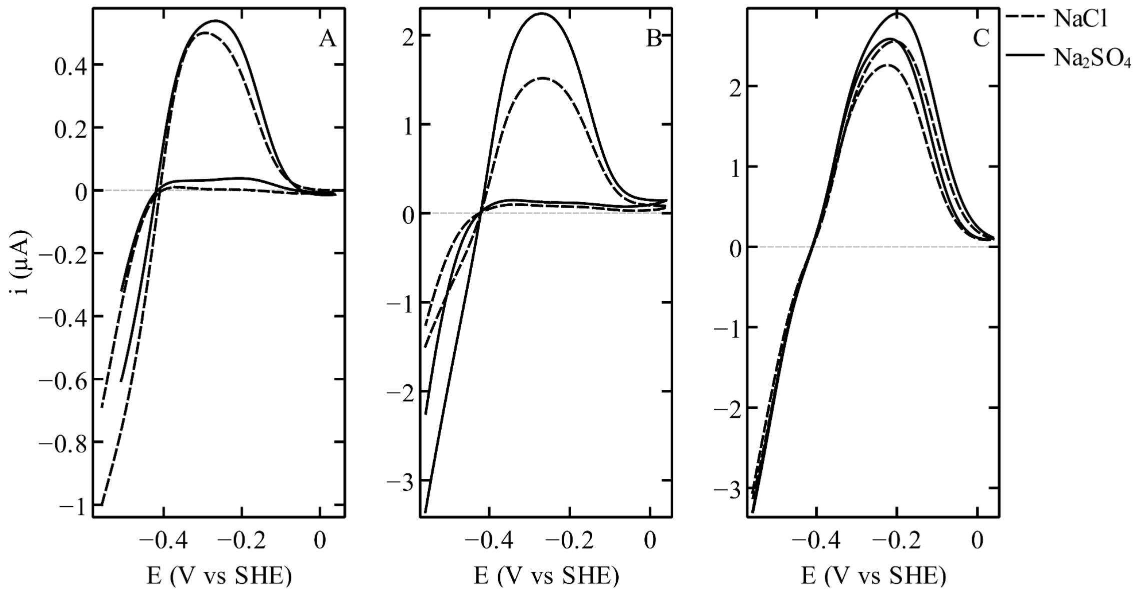


Figure S2: Effect of chloride inhibition on CpIII WT (panel A), MeII WT (panel B), CpIII ΔC221 (panel C). In solid line voltammograms recorded with 0.1M $Na_2SO_4$, in dash line voltammograms recorded with 0.12M NaCl. Other conditions: pH 7, 30°C, 3000 rpm.

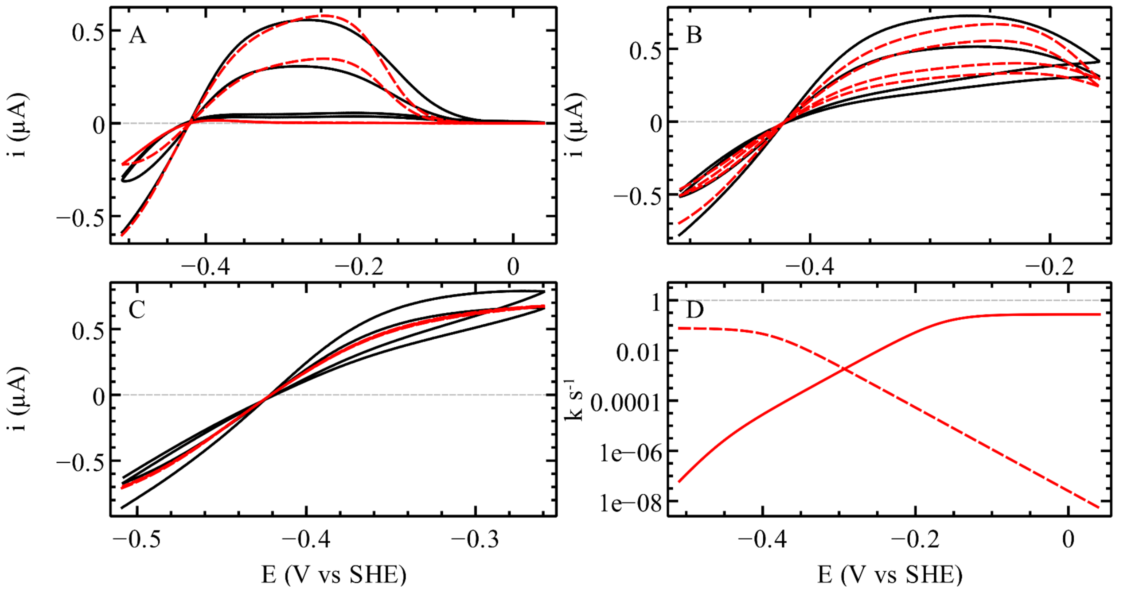


Figure S3: fit of the 1st model to three selected voltammograms of WT CpIII among those of figure 4B in the main text. Panels A, B and C show the experimental data in black and the fits of the 1st model in dash red lines. Panel D shows the potential dependence of $k_i$ (solid line) and $k_a$ (dash line) obtained from the global fit.

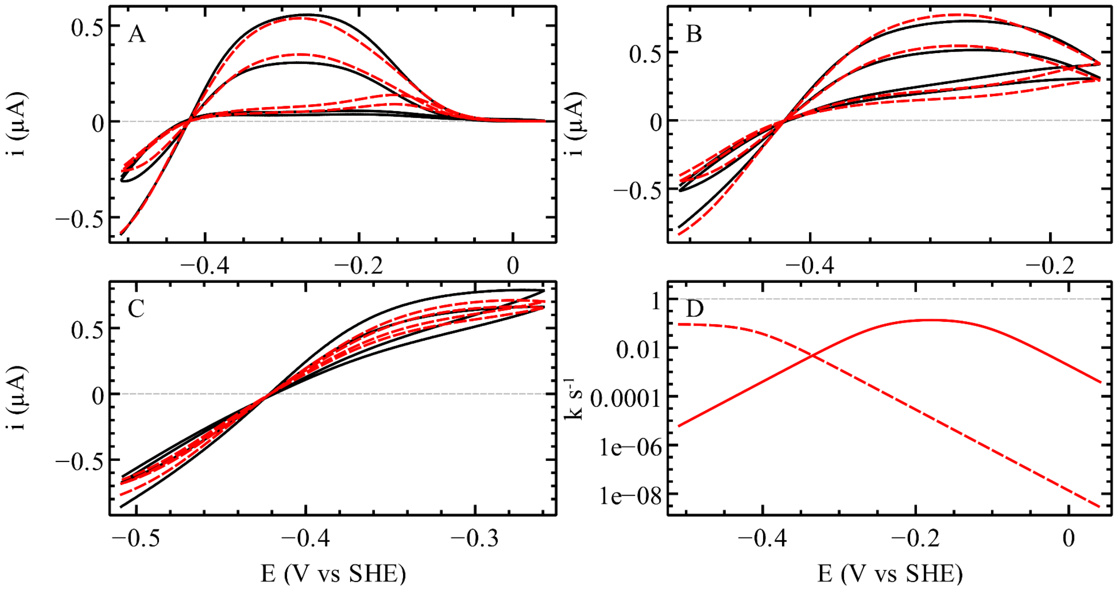


Figure S4: fit of the 2nd model to three selected voltammograms of WT CpIII among those of figure 4B in the main text. Panels A, B and C show the experimental data in black and the fits of the 2nd model in dash red lines. Panel D shows the potential dependence of $k_i$ (solid line) and $k_a$ (dash line) obtained from the global fit.

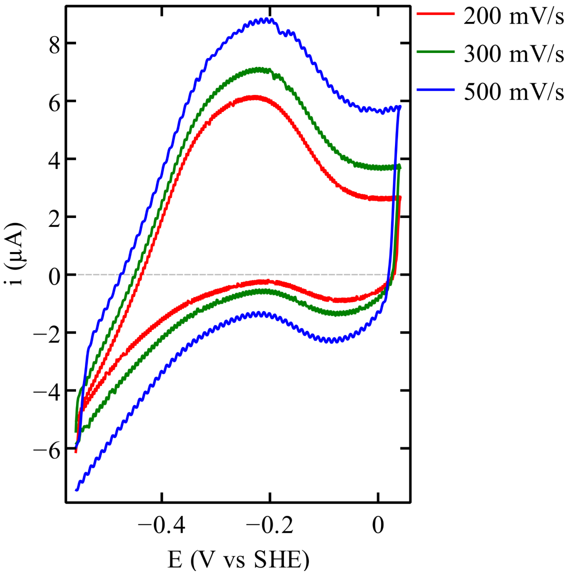


Figure S5: CVs of CpIII WT at high scan rate: 200 mV/s in red; 300 mV/s in green; 500 mV/s in blue. Other conditions: pH 7; 30C; 3000 rpm.

### Structural considerations

#### Sequence alignments

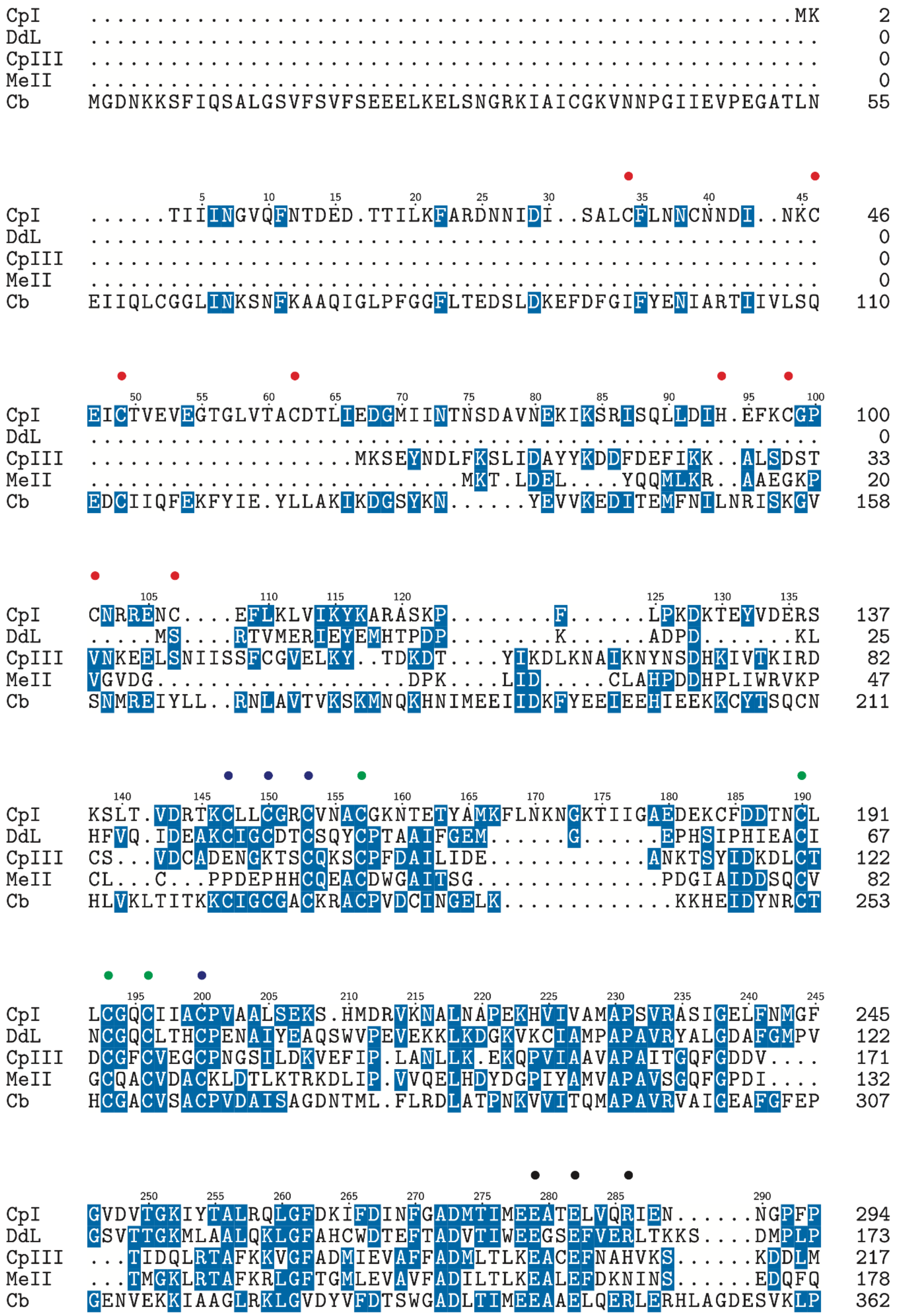

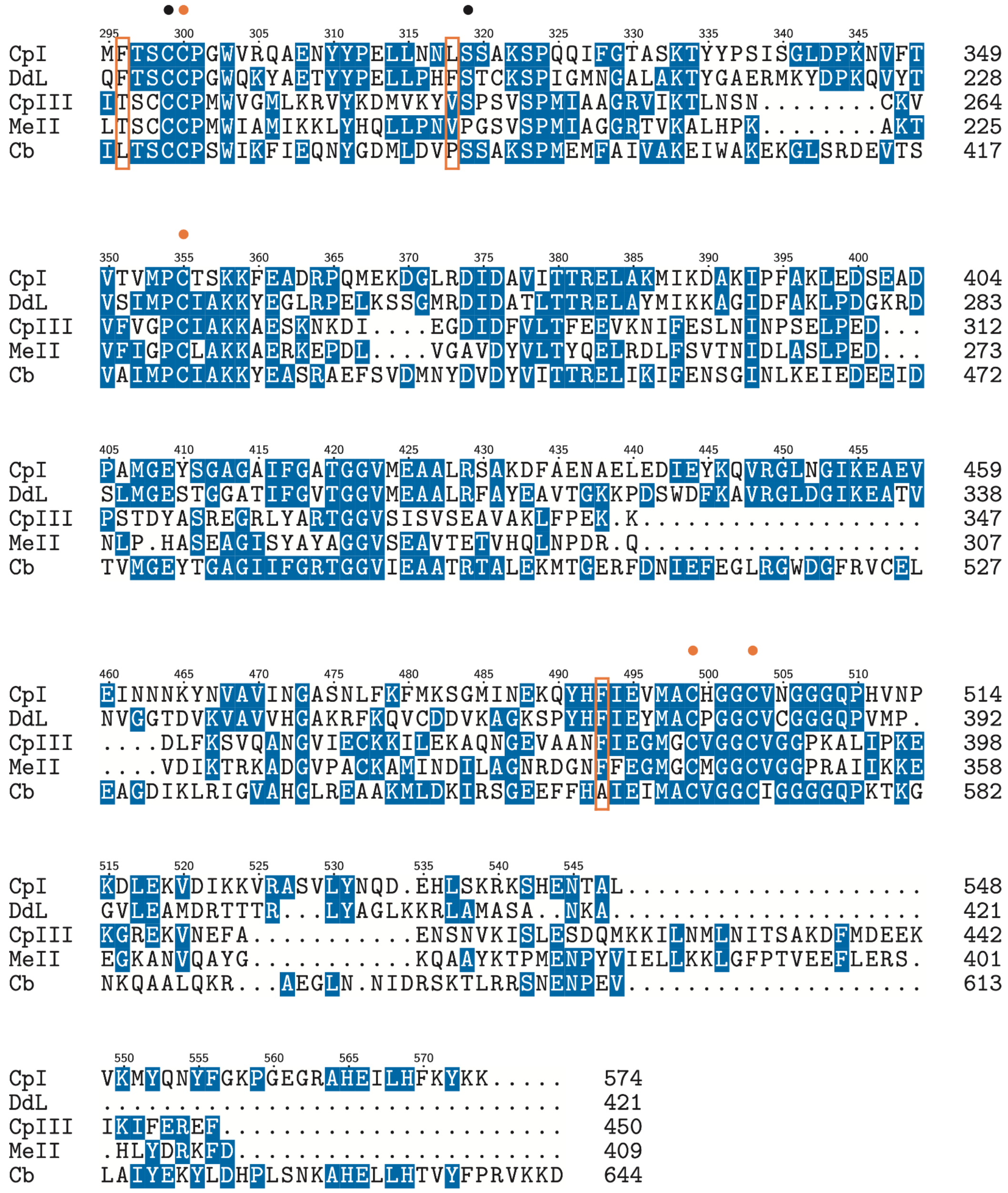


Figure S6. ClustalO alignment of the sequences of CpI, *D. desulfuricans* (Dd, Large subunit), CpIII, MeII and Cb FeFe hydrogenases. The residues involved in the binding of the two N-terminal clusters of CpI are marked by red dots. Those involved in the binding of the two accessory clusters in Dd, CpIII and MeII hydrogenases are marked by blue and green dots (note the peculiar 1st run of cysteines in CpIII and MeII). The residues that bind the cubane of the H cluster are marked in orange. Those that are part of the conserved proton transfer chain in group A hydrogenases are marked by black dots. The Cb L364, P386 and A561 residues, identified in ref [6], are framed by orange boxes.

#### Alphafold calculations

The model structures were calculated with ColabFold v1.5.2, AlphaFold2 using MMseqs2, relaxing the predicted structures using amber force fields[7].

#### The coordination of the accessory clusters in CpIII

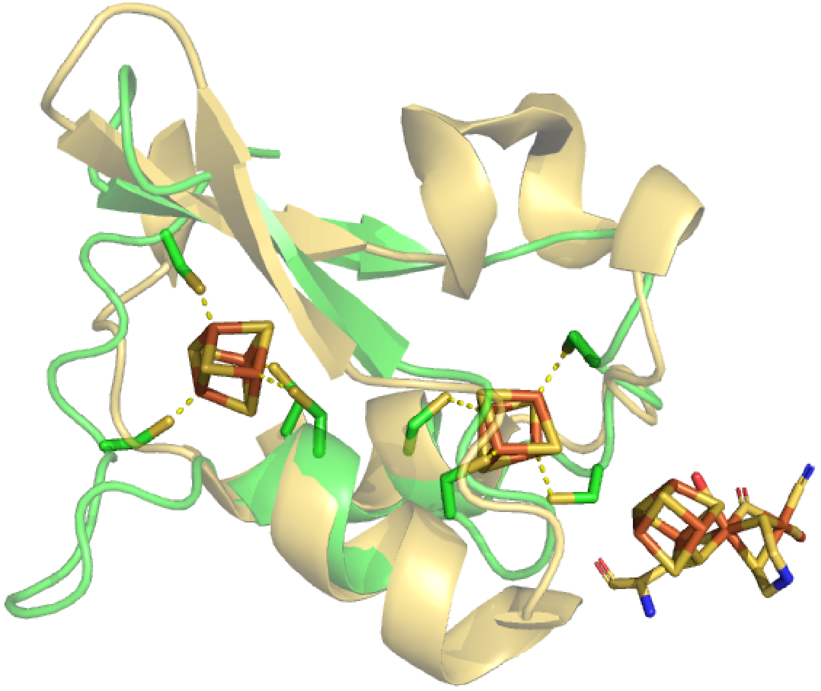

Figure S7. The environment of the two accessory [4Fe4S] clusters of CpIII. The alphafold structure of CpIII (green) is aligned to that of CpI (pdb 6N59). The CpIII first run of cysteines (leftmost), coordinating the distal [4Fe4S] cluster, is $C^{83}x_2Cx_9C^{96}$, instead of $C^{147}x_2Cx_2C^{153}$ in CpI (blue dots in SI fig S6). The loop between C86 and C96 in CpIII is seen here on the left. This motif is highly variable in group B hydrogenases[8,9].
